# Tuba1a is uniquely important for axon guidance through midline commissural structures

**DOI:** 10.1101/2020.05.05.079376

**Authors:** Georgia Buscaglia, Jayne Aiken, Katelyn J. Hoff, Kyle R. Northington, Emily A. Bates

## Abstract

Developing neurons undergo dramatic morphological changes to appropriately migrate and extend axons to make synaptic connections. The microtubule cytoskeleton, made of α/β-tubulin dimers, drives neurite outgrowth, promotes neuronal growth cone responses, and facilitates intracellular transport of critical cargoes during neurodevelopment. *TUBA1A* constitutes the majority of α-tubulin in the developing brain and mutations to *TUBA1A* in humans cause severe brain malformations accompanied by varying neurological defects, collectively termed tubulinopathies. Studies of *TUBA1A* function *in vivo* have been limited by the presence of multiple genes encoding highly similar tubulin proteins, which prevents TUBA1A-specific antibody generation and makes genetic manipulation challenging. Here we present a novel tagging method for studying and manipulating *TUBA1A* in cells without impairing tubulin function. Using this tool, we show that a *TUBA1A* loss-of-function mutation *TUBA1A^N102D^* (*TUBA1A^ND^*), reduced the amount of TUBA1A protein and prevented incorporation of TUBA1A into microtubule polymers. Reduced Tuba1a α-tubulin in heterozygous *Tuba1a^ND/+^* mice significantly impacted axon extension and impaired formation of forebrain commissures. Neurons with reduced Tuba1a caused by *Tuba1a^ND^* had altered microtubule dynamics and slower neuron outgrowth compared to controls. Neurons deficient in Tuba1a failed to localize microtubule associated protein-1b (Map1b) to the developing growth cone, likely impacting reception of developmental guidance cues. Overall, we show that reduced Tuba1a is sufficient to support neuronal migration, but not axon guidance, and provide mechanistic insight as to how *TUBA1A* tunes microtubule function to support neurodevelopment.

## Introduction

Mammalian brain development is a complex process that requires precise coordination of multiple cell types and extracellular cues to form a fully specified tissue. Despite many advances in understanding the cellular and molecular players involved in brain development, there is still much that remains unknown. Insights into the molecular pathways governing neurodevelopment can be gained from studying genetic mutations that disrupt specific aspects of brain development. Severe cortical and neurodevelopmental phenotypes associated with mutations that disrupt tubulin genes, termed tubulinopathies, have recently been described in humans [1–4]. Tubulinopathy mutations cause a spectrum of neurodevelopmental phenotypes, but frequently involve cortical malformations such as lissencephaly, agenesis or hypoplasia of the corpus callosum, and cerebellar hypoplasia [1, 2, 4, 5]. Recent studies of human tubulinopathy mutations have revealed that each variant may impact different aspects of microtubule function, such as protein folding, polymerization competency, and microtubule-associated protein (MAP)-binding, among others [6–9]. Tubulin mutations can therefore be used to interrogate the requirement for specific aspects of microtubule function throughout neurodevelopment.

Developing neurons must migrate to the correct location, extend axons to meet sometimes distant synaptic partners and form functional connections. Throughout this process, neurons undergo dramatic morphological changes that require coordinated interaction between the cytoskeleton and the extracellular environment. In post-mitotic neurons, microtubule polymers made of α/β-tubulin dimers facilitate nucleokinesis and cellular migration, support growth cone navigation, promote axon formation and form the tracks upon which intracellular trafficking occurs [10–13]. The microtubule network needs to be precisely controlled to fulfill diverse functions in neurons. Microtubule properties can be modulated through post-translational modifications (PTMs), association with MAPs and through the particular tubulin genes, or isotypes, that a cell expresses [14]. The human genome contains at least nine unique α- and ten unique β-tubulin genes [15, 16]. The α-tubulin isotype encoded by the gene *TUBA1A* is abundant in the brain and is the most highly expressed α-tubulin in post-mitotic, developing neurons [17–19]. *TUBA1A* mutations are highly represented in cases of human tubulinopathies [20], suggesting that *TUBA1A* plays an important role in neurodevelopment. However, due to the high degree of sequence conservation between α-tubulin genes, it has been historically challenging to study *TUBA1A* function *in vivo*, due to the limited availability of tools.

Mouse models harboring mutations to *Tuba1a* can be used as tools to interrogate the function of Tuba1a in the context of the neuronal milieu. As tubulin genes are often required for life and the nucleotide sequence between isotypes is conserved, generation of mutant mouse lines to study *Tuba1a* function *in vivo* has been challenging. To date, only a handful of *Tuba1a* mutant mouse lines have been generated, three by ENU-induced forward genetic screens and one by site-directed CRISPR gene editing [21–23]. We previously showed that the ENU-induced *Tuba1a^N102D^* allele (*Tuba1a^ND^*) impaired microtubule function in both *S. cerevisiae* and mice [22]. Homozygous *Tuba1a^ND^* mice exhibit severely impaired brain development and are neonatal lethal, similar to phenotypes seen in the *Tuba1a^null^* and *Tuba1a*-R215* mutant mice [20–22]. In homozygous *Tuba1a^ND^*, *Tuba1a^Null^* and *Tuba1a*-R215* mice, as well as many *TUBA1A* tubulinopathy patients, cortical migration and commissural formation are severely disrupted, making it difficult to infer whether axon pathfinding is a direct consequence of altered Tuba1a function or if it is secondary to abnormal cortical layering and migration. *Tuba1a^ND/+^* heterozygous mutant mice have reduced Tuba1a function during brain development, which was sufficient to support survival and cortical migration resulting in normal cortical layers [22, 24], but does not support formation of commissures. Therefore, *Tuba1a^ND/+^* heterozygous animals can provide insight into how Tuba1a contributes specifically to axon pathfinding.

Here, we show that a reduction in developmental Tuba1a protein causes specific impairments in axon guidance through large brain commissures. Using a novel tubulin visualization technique, we demonstrate that the *TUBA1A^N102D^* mutation prevents incorporation of *TUBA1A* into microtubule polymers in cells. In mice, heterozygous *Tuba1a^ND/+^* brains fail to form the corpus callosum, anterior and hippocampal commissures due to apparent deficits in axon pathfinding through the midline. Cultured neurons from *Tuba1a^ND/+^* and wild-type cortices revealed that *Tuba1a^ND/+^* neurons have shorter neurites than wild-type and grow slower overall, potentially due to alterations in microtubule polymerization dynamics. Further, we demonstrate that *Tuba1a^ND/+^* neurons fail to localize at least one critical developmental MAP to the developing growth cone. Collectively, our data present evidence for precise mechanisms by which loss of Tuba1a causes axonal pathfinding deficits during development.

## Materials and Methods

### Animals

All animal research was performed in accordance with the Institutional Animal Care and Use Committee at the University of Colorado School of Medicine. All mice used were maintained on a 129S1/C57Bl6 genetic background. Mice were kept on a 12:12 light:dark cycle with *ad libitum* access to food and water. *Tuba1a^ND^* and wild-type littermate mice were maintained on water supplemented with 0.2g/L MgSO_4_ to promote *Tuba1a^ND/+^* survival and ability to reproduce. For timed mating male and female mice were introduced overnight and separated upon confirmation of mating, which was then considered embryonic day 0.5. Male and female mice were represented in all studies. All mice were genotyped by PCR amplification of tail DNA followed by Sanger sequencing to differentiate homozygous or heterozygous *Tuba1a^ND/+^* mice from wild-type. Primers used to amplify mouse DNA for genotyping were: forward:TGGATGGTACGCTTGGTCTT; reverse: CTTTGCAGATGAAGTTCGCA; and sequencing: GTCGAGGTTTCTACGACAGATATC.

### Histology

Mice were anesthetized and trans-cardially perfused with 0.1M NaCl and 4% paraformaldehyde (PFA) for histology. Tissues of interest were dissected and post-fixed in 4% PFA. Tissue sectioning was performed on a CM1520 cryostat (Leica, Wetzlar, Germany) and 30μm cryosections were obtained for all histology. For luxol fast blue staining, sections from brain were stained using a 0.1% luxol fast blue solution. For immunofluorescence studies PFA-fixed tissues or cells were blocked in phosphate-buffered saline (PBS) containing 5% goat serum or bovine serum albumin (BSA) with 0.3% Triton-X 100. Primary and secondary antibodies were diluted in PBS containing 1% BSA with 0.1% Triton-X 100.

### Electron Microscopy

Mice used for electron microscopy were perfused with 0.1M NaCl and 2.5% glutaraldehyde 4% PFA, after which the brain was dissected and post-fixed in 2.5% glutaraldehyde 4% PFA overnight at 4°C. Following post-fixation, brains were sent for sectioning and imaging by the CU School of Medicine Electron Microscopy Core facility.

### Plasmids and Reagents

The hexahisitidine (His6) epitope tag was inserted in the α-tubulin internal loop region [25–27]. Codon optimization for *rattus norvegicus* (https://www.idtdna.com/codonopt) was used to generate the His6 sequence CATCACCACCATCATCAC, which was inserted into the coding region of human *TUBA1A* from the Mammalian Genome Collection (clone ID:LIFESEQ6302156) between residues I42 and G43. Gibson cloning was used to insert the gBlock of *TUBA1A* internally tagged with His6 (*TUBA1A*-His6) into the pCIG2 plasmid (shared by Dr. Matthew Kennedy, University of Colorado) linearized with NruI and HindIII. *TUBA1A*-His6 incorporation was confirmed by sequencing across the complete *TUBA1A* coding region. The *TUBA1A^T349E^* (*TUBA1A^TE^)* polymerization incompetent, and *TUBA1A^E255A^* (*TUBA1A^EA^*) highly polymer-stable α-tubulin mutants were identified and described in prior publications [28–30].The GFP-MACF43 vector was shared by Dr. Laura Anne Lowery (Boston College) and Dr. Casper Hoogenraad (Utrecht University). Myr-TdTomato plasmid DNA was shared from Mark Gutierrez and Dr. Santos Franco (University of Colorado).

### Cell Culture and Nucleofection

Cos-7 cells (Thermo Fisher Scientific, Waltham, MA; ATCC^®^ CRL-1651™) were cultured in a 37°C humidified incubator with 5% CO_2_ in DMEM (Gibco) supplemented with 10% fetal bovine serum (Gibco), 1mM sodium pyruvate (Thermo), and penn/strep (1,000 IU/ 1,000 μg/mL; Thermo). Cos-7 cells were transfected with 2.5μg of hexahistidine (His6) tagged *TUBA1A* plasmid DNA using Lipofectamine 3000 (Invitrogen). For Cos-7 proteasome inhibition assays, 5μm Lactacystin A [31, 32] was added to normal culture medium for 24 hours, the day following transfection with *TUBA1A*-His6 constructs. Dissociated neurons were cultured from male and female P0-P2 mouse or rat cortices. Brains were removed and placed into HBSS (Life Technologies) supplemented with 1M HEPES (Life Technologies) and 1mM kynurenic acid (Tocris Bioscience, Bristol, UK). Meninges were removed and cortices were dissected and cut into approximately 1mm pieces. Cortical pieces were triturated to a single-cell suspension using glass Pasteur pipettes. Cortical neurons were plated onto 35mm Poly-D-Lysine coated glass-bottom culture dishes at a density of 350,000 cells (Willco Wells, HBSt-3522). For nucelofected mouse and rat neurons, 4 μg of plasmid DNA was introduced to 4×10^6^ neurons using an AMAXA nucleofection kit (VPG-1001, VPG-1003; Lonza). AMAXA-nucleofected cells were plated in 35mm glass bottom imaging dishes. Neurons were maintained in a 37°C humidified incubator with 5% CO_2_ in phenol-free Neurobasal-A medium (Life Technologies) supplemented with B-27 (Thermo Fisher Scientific, Waltham, MA), Penn/strep (Thermo), GlutaMax (Thermo), 5ng/mL β-FGF (Gibco), and Sodium Pyruvate (Thermo).

### RNA isolation + RTPCR

RNA was isolated from Cos-7 cells, 48-hours post-transfection using TRIzol Reagent (Thermo; 15596026). RNA concentration and purity were determined using a spectrophotometer, then cDNA was synthesized using the RT2 First Strand Kit (Qiagen, Hilden, Germany; 330401). qRT-PCR reactions were prepared with SYBR Green RT-PCR Master mix (Thermo; S9194) and run with a CFX Connect Real-Time System (Bio-Rad). Samples were run in triplicate, results were analyzed in Excel. All qPCR data presented in this manuscript was normalized to expression of GFP, which was present on the same plasmid as *TUBA1A*-His6 constructs. Wild-type *TUBA1A* mRNA quantity was set to = 1 and *TUBA1A^ND^* relative mRNA quantity was presented relative to wild-type. For all qRT-PCR experiments 3 biological replicates were used per genotype.

### Neuron Immunocytochemistry

DIV 3 primary cortical neurons were washed with PBS and fixed with a fixation solution of 4% PFA and 0.2% glutaraldehyde in PBS for 15 minutes at room temperature. For tubulin extraction, cells were washed with PBS followed by PHEM buffer, then soluble tubulin dimers were extracted using 0.1% triton with 10μM taxol and 0.1% DMSO in PHEM buffer. Extracted cells were fixed with 2% PFA and 0.05% glutaraldehyde in PBS for 10 minutes, washed with PBS and then reduced in 0.1% NaBH4 in PBS for 7 minutes at room temperature. Cells were then washed with PBS and blocked in 3% BSA and 0.2% Triton in PBS for 20 minutes at room temperature, with agitation. Immunostaining was performed using primary antibodies directed against: 6X-Histidine (Invitrogen, 4A12E4 37-2900; 1:500), total α-tubulin (Sigma, DM1A T6199; 1:5,000), Acetylated Tubulin (Sigma, T7451; 1:1,000), Tyrosinated Tubulin (Chemicon, MAB1864; 1:1,000), Map1b (Santa Cruz Biotech, sc-135978; 1:500), Map2 (Novus Biologicals, NB300-213; 1:2,000). Primary antibodies were diluted in blocking buffer and incubated overnight at 4°C in a humidified chamber. After primary antibody staining, cells were washed three times with PBS. Fluorescently-conjugated secondary antibodies were diluted 1:500 in blocking buffer and incubated for 2 hours at room temperature, protected from light. Secondary antibodies were from Life Technologies (Carlsbad, CA) all used at 1:500. Alexa Fluor 568-conjugated Phalloidin (Thermo, A12380; 1:20) was added during secondary antibody incubation for labeling of filamentous actin. Tissues or cells of interest were mounted onto glass microscope slides and sealed with glass coverslips and aqueous mounting media containing DAPI (Vector Laboratories, #H-1200) and imaged on a Zeiss 780 or 880 confocal microscope with a 40X or 63X oil objective.

### Microtubule Dynamics and Neuron Growth Rates

Primary neurons from wild-type and *Tuba1a^ND/+^* neonatal mouse cortices were cultured as described above. Prior to plating, mouse cortical neurons were nucleofected with 4μg each of GFP-MACF43 and Myr-TdTomato plasmid DNA. Following 1 day in culture, neurons were transferred to a 37°C imaging chamber and time-lapse images of GFP-MACF43 comets and Myr-TdTomato membrane label were acquired using a Zeiss 780 confocal microscope with 40X oil objective. Following acquisition of baseline images, images were acquired every 2 seconds for 2 minutes. A total of four time-lapse videos were acquired per neuron, with a 10-minute waiting period in between imaging sessions. Data from all four GFP-MACF43 time points were pooled by cell to generate a cell average and then grouped by genotype for further analysis. Membrane-bound Myr-TdTomato images were acquired at the beginning and end of the imaging period and were used to track neuronal growth. Kymographs of GFP-MACF43 comets in single neurites were generated from GFP-MACF43 videos in ImageJ/FIJI (National Institutes of Health) and were used to assess microtubule polymerization velocity, along with the duration and distance of individual plus-end polymerization events. In brief, polymerization velocity was determined by measuring the change in position (X_2_-X_1;_ μm) divided by the change in time (Y_2_-Y_1;_ min) for each GFP-MACF43 comet, duration assessed the total time (Y_2_-Y_1_) each GFP-MACF43 comet spent moving in seconds, and distance assessed the total distance (X_2_-X_1_) covered by each polymerization event in μm.

### Western Blotting

Protein was isolated from brains of P0-P2 mice by dounce homogenization and ultra-centrifugation. Tubulin-enriched protein fractions with MAPs were isolated according to a previously established protocol [33]. Cos-7 cell protein was extracted using a Tris-Triton lysis buffer with protease inhibitor cocktail (Sigma). Protein concentrations were assessed using a BCA assay (Thermo), and relative concentration was determined using a Synergy H1 microplate reader (BioTek Instruments, Winooski, VT). 5μg of either whole brain lysate or tubulin-enriched protein fraction was loaded per lane and run on 4-20% Mini-Protean TGX Stain-Free precast gels (4568093; Bio-Rad Laboratories, Hercules, CA) at 150mV for 1hr. Prior to protein transfer, Stain-Free gels were activated with UV light for 1 minute and imaged for total protein on gel using a ChemiDoc MP imager (Bio-Rad). Proteins were transferred to PVDF blotting membranes (Bio-Rad) in standard 25mM Tris-base, 192mM glycine, 15% methanol transfer buffer, or transfer buffer optimized for high molecular-weight proteins (>200kDa) by the addition of 0.025% SDS. Blots were transferred at 4°C and 75V for either 1 hour for standard molecular-weight proteins, or 3 hours for high molecular-weight proteins. Immediately following transfer, PVDF membranes were rinsed in TBST and imaged to quantify the total protein on blot using UV-activated Stain-Free blots. Gels were also imaged post-transfer to assess transfer efficiency for each blot. Membranes were blocked in Tris-buffered Saline containing 0.1% Tween-20 (TBST) with 5% BSA for 1 hour and incubated in primary antibody diluted in TBST containing 1% BSA overnight at 4°C. Primary antibodies were: mouse anti 6X-Histidine (Invitrogen, 4A12E4; 1:500), chicken anti-GFP (Invitrogen, A10262; 1:1,000), and mouse anti-Map1b (Santa Cruz, sc-135978; 1:500). Blots were incubated in HRP-conjugated Goat-anti-mouse (1:5,000; Santa Cruz) secondary antibody diluted in TBST containing 0.5% BSA with streptavidin-HRP (Bio-Rad, 1:10,000) for 1 hour at room temperature. Blots were developed in ECL solution for 5 minutes at room temperature (Bio-Rad) and imaged.

### Experimental design and statistical analyses

Band volume of all Western blots was analyzed using Image Lab software (Bio-Rad). Fluorescence intensity measurements, area and morphological assessment, kymograph generation, and quantification of EM images was performed using ImageJ/FIJI software (NIH) and Excel (Microsoft). Statistical analyses were performed, and graphs were created using Prism version 8.0 (GraphPad). Most graphs display all data points to accurately represent the variability in each dataset, except in cases where such presentation obscured visibility. For all statistical analyses, statistical significance was considered to be p < 0.05. Statistical analyses used in each experiment are indicated in their respective figure legends. For all graphs mean ± SEM was reported unless otherwise noted. Normality of each dataset was assessed using a Shapiro-Wilk test. In datasets with two groups, parametric data was analyzed using a Student’s t-test, while non-parametric data was assessed by Mann-Whitney *U* analysis of medians. Multiple groups were compared by one-way or two-way ANOVA and analyzed *post hoc* by either a Bonferroni or Kruskal-Wallis test for parametric and non-parametric data, respectively.

## Results

### *Tuba1a* is required for formation of midline commissural structures

*TUBA1A* is a major component of developing neuronal microtubules, and is critical for proper brain development [20]. Human *TUBA1A*-tubulinopathy patients and homozygous *Tuba1a* mutant mouse models both exhibit severe brain malformations. *Tuba1a^ND/+^* heterozygous mutant mice undergo normal cortical migration, display comparable brain weight to wild-type littermates at birth, and survive to adulthood [22, 24]. However, *Tuba1a^ND/+^* brains exhibit agenesis of the corpus callosum and abnormal formation of the anterior and hippocampal commissures (Fig. 1A). In wild-type mice, nascent callosal ‘pioneer’ axons extend through midline at E15.5, and early ‘follower’ axons begin extending at E17 in mice [34]. Evidence of abnormal callosal projections were apparent as early as P0 in *Tuba1a^ND/+^* brains, as seen by the early formation of aberrant axon bundles adjacent to the callosum, known as Probst bundles (Fig. 1A)[35]. In addition to agenesis of the corpus callosum at midline, lateral regions of adult *Tuba1a^ND/+^* corpus callosum were found to be significantly thinner than wild-type (Fig. 1B). Similarly, adult *Tuba1a^ND/+^* anterior commissures are smaller than that of wild-type littermates (Fig. 1C). In wild-type mice, corpus callosum thickness and anterior commissure area both increased significantly between P0 and adulthood; however, normal postnatal expansion of these tracts was not evident in *Tuba1a^ND/+^* mice (Fig. 1B, C). *Tuba1a^ND/+^* axons fail to organize into typical midline commissural structures, indicating that half of the normal complement of Tuba1a during brain development is not sufficient for commissural axon guidance.

**Figure 1.**
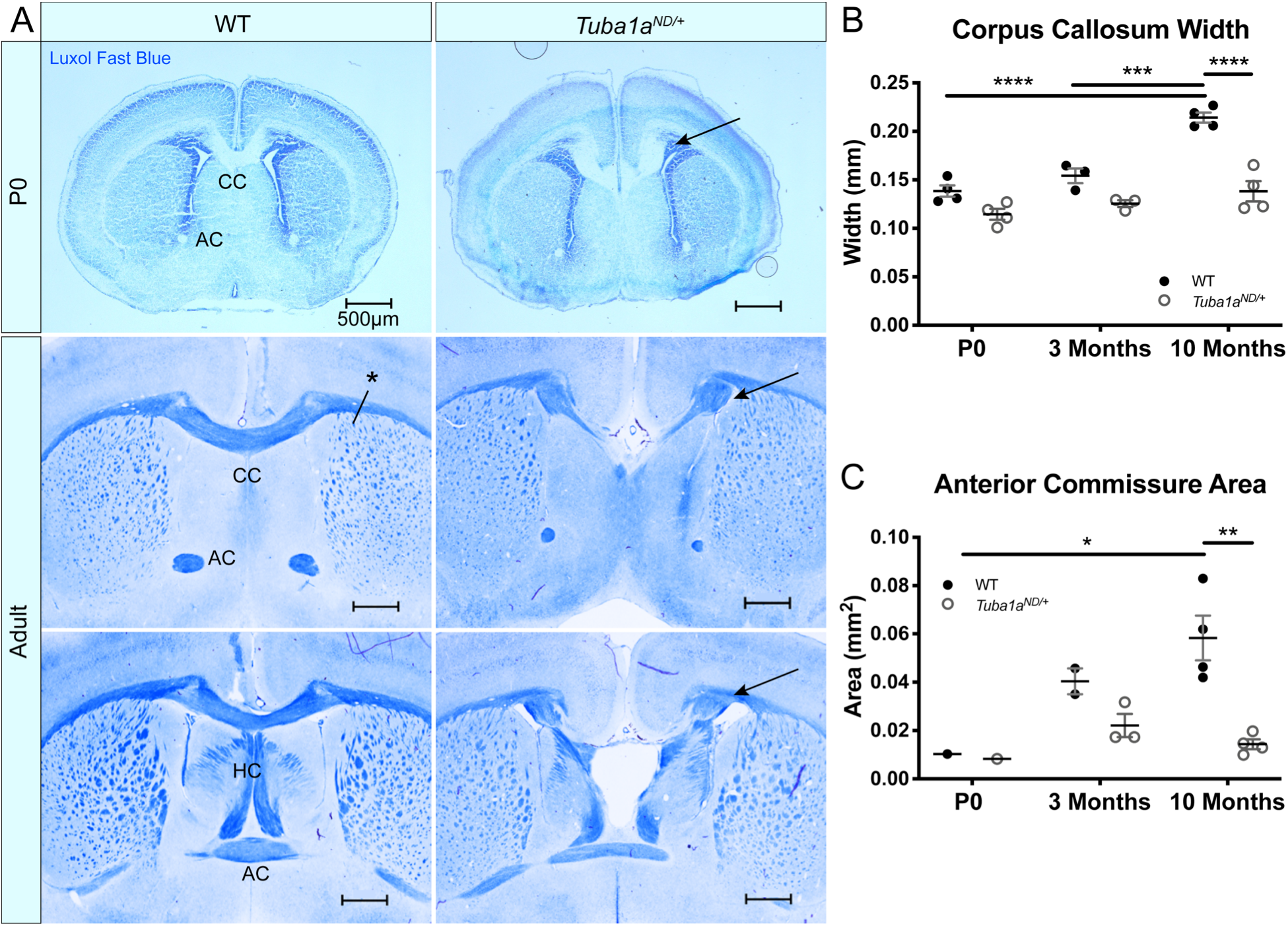
Tuba1a is required for formation of midline commissural structures. ***A.*** Luxol fast blue-stained coronal brain sections from postnatal day 0 (P0; top) and Adult (middle-bottom) wild-type and *Tuba1a^ND/+^* mice. Images portray abnormal midline commissural formation in *Tuba1a*^*ND/+*^ mouse brains, with labels highlighting the corpus callosum (CC), anterior commissures (AC), and hippocampal commissure (HC). Scale bars are 500μm. ***B.*** Scatter plot representing corpus callosum width at P0, 3-months, and 10-months-old. Arrows indicate Probst bundles. Asterisk in *A.* shows where measurements for *B.* were obtained. ***C.*** Scatter plot displaying anterior commissure area in P0, 3-month, and 10-month-old wild-type and *Tuba1a*^*ND/+*^ brains. Wild-type and *Tuba1a*^*ND/+*^ measurements compared by two-way ANOVA. * p<0.05; ** p<0.01; *** p<0.001; and **** p<0.0001.

Examination of sagittal brain sections taken directly at midline revealed dramatic disorganization of corpus callosum axons in *Tuba1a^ND/+^* brains (Fig. 2A). Compared to wild-type, *Tuba1a^ND/+^* midline commissural axons were largely absent, and the existing axons failed to organize into a tract with uniform orientation (Fig. 2A). Despite dramatic differences between wild-type and *Tuba1a^ND/+^* callosal axon organization, *Tuba1a^ND/+^* axons were highly colocalized with immunolabeled myelin sheaths (Fig. 2A). To further assess the impact of *Tuba1a^ND/+^* substitution on callosal axon morphology and myelination, we performed electron microscopy (EM) in both wild-type and *Tuba1a^ND/+^* corpus callosi. Due to the scarcity of axons present directly at midline in the *Tuba1a^ND/+^* corpus callosum, we sampled a region of corpus callosum 2mm lateral to midline for both wild-type and *Tuba1a^ND/+^* animals (Fig. 2B). EM images revealed a striking difference in axon density between wild-type and *Tuba1a^ND/+^* corpus callosi (Fig. 2B, D; p=0.03). Myelin thickness, measured by G-ratio, was similar between wild-type and *Tuba1a^ND/+^* brains (Fig. 2C; p=0.34), as was axon diameter (Fig. 2E; p=0.14). There was a trend towards decreased myelination in *Tuba1a^ND/+^* animals (p=0.07), but this difference was not statistically significant (Fig. 2F). These data provide evidence that *Tuba1a^ND/+^* callosal axons do not correctly organize to form a commissure. Previously published data indicated that reduced developmental Tuba1a function is tolerable for cortical neuron migration [22]; however, our results indicate that reduced Tuba1a is not sufficient to support axon pathfinding.

**Figure 2.**
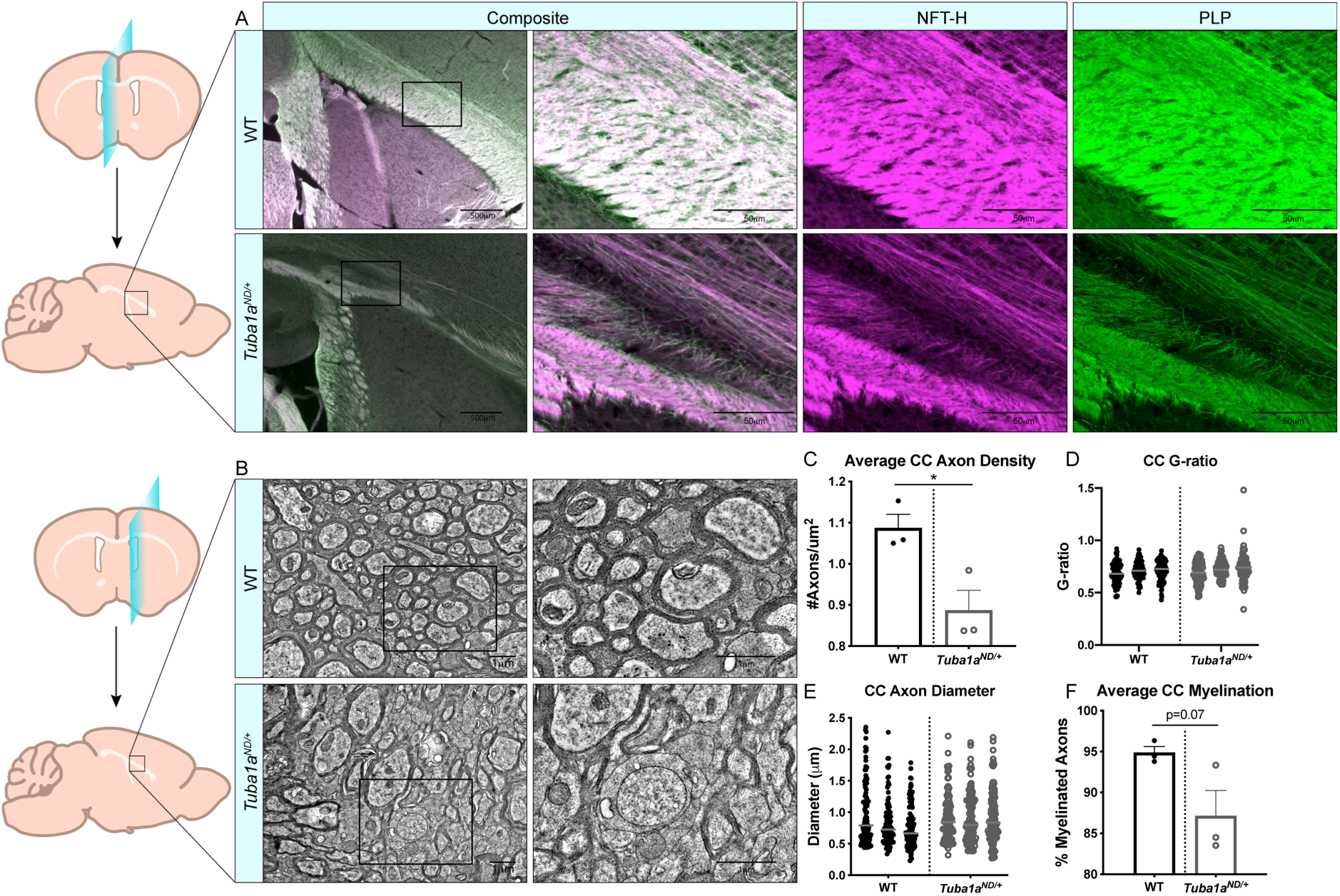
Axon density is reduced by Tuba1a^ND/+^ in midline corpus callosum. ***A.*** Sagittal brain sections at midline from wild-type (top) and *Tuba1a*^*ND/+*^ (bottom) brains stained with neurofilament-heavy (NFT-H; magenta) to label axons, and proteolipid protein (PLP; green) to label myelin. Images were taken at 4X (left) and 20X (right) magnification with 500μm and 50μm scale bars, respectively. ***B.*** Electron microscopy (EM) images of sagittal wild-type (top) and *Tuba1a*^*ND/+*^ (bottom) sections of corpus callosum 2mm adjacent to midline. Enlarged regions on right are denoted by boxes, scale bar is 1μm. ***C.*** Scatter plot of G-ratio measurements for wild-type and *Tuba1a*^*ND/+*^ axons. Data points represent individual myelinated axons and are clustered by animal (N=3 mice, n=100 axons; p=0.34). ***D.*** Scatter plot representing axon density of wild-type and *Tuba1a*^*ND/+*^ axons in analyzed EM images. Data points represent average values for each animal (n=6 regions containing same total area; p=0.03). ***E.*** Scatter plot of axon diameters in wild-type and *Tuba1a*^*ND/+*^ corpus callosum by EM. Only those axons captured in cross section were assessed for diameter, as skewed axons provide inaccurate measurements. Data points represent individual axons and are clustered by animal (n=100; p=0.14). ***F.*** Scatter plot representing the average percent of myelinated axons per animal in wild-type and *Tuba1a*^*ND/+*^ corpus callosum. 6 regions containing the same total area were assessed (n=6; p=0.07). Statistical differences between means of wild-type and *Tuba1a*^*ND/+*^ datasets were assessed by t test, with * p<0.05.

### *TUBA1A^ND^* α-tubulin does not incorporate into neuronal microtubules

Reduced developmental Tuba1a is not adequate to support long-distance axon targeting, but the molecular and cellular mechanisms by which Tuba1a alters neuronal microtubule function are unknown. The *Tuba1a^ND^* allele was previously demonstrated to cause partial loss of microtubule function in yeast and mice [22, 24]. However, the mechanism by which *Tuba1a^ND^* substitution leads to loss-of-function (LOF) is unclear. The neuronal microtubule network is complex, containing many different tubulin isotype proteins, PTMs, and MAPs decorating the lattice, all of which can modify functional microtubule properties. Further, the high degree of sequence similarity between α-tubulin isotypes has precluded study of individual tubulin isotypes at the protein level *in vivo*. No α-tubulin isotype or species-specific antibodies exist for TUBA1A, and prior attempts to tag TUBA1A neuronal microtubules with N- or C-terminal fusion proteins have had detrimental effects on protein function [36]. These challenges have made the direct visualization of specific tubulin isotypes or mutant tubulin proteins in neurons historically difficult. Thus, the ways in which TUBA1A specifically contributes to neuronal microtubule protein function have been difficult to ascertain. To address the need for better tools to study TUBA1A protein, we generated a functional hexahistidine (His6)-tagged TUBA1A construct based on a previously identified internal loop in the α-tubulin protein for *in vivo* visualization and manipulation of microtubules [26] (Fig. 3A). We inserted a His6 tag into an internal loop of TUBA1A between residues I42 and G43. This region of α-tubulin was previously shown to tolerate addition of amino acids without disrupting tubulin function [26], and previous groups added a His6 tag in this loop to affinity purify tubulin subunits for reconstituted systems [25, 27]. However, to our knowledge this internal His6 tag on α-tubulin has never been used for immunohistochemical assays to visualize the microtubule network *in vivo*. Ectopically expressed wild-type *TUBA1A*-His6 is incorporated into Cos-7 cell microtubules (Fig. 3B). Compared to traditional immunolabeling of cellular microtubules (DM1A), *TUBA1A*-His6 provides excellent labeling of microtubule polymers (Fig. 3B).

**Figure 3.**
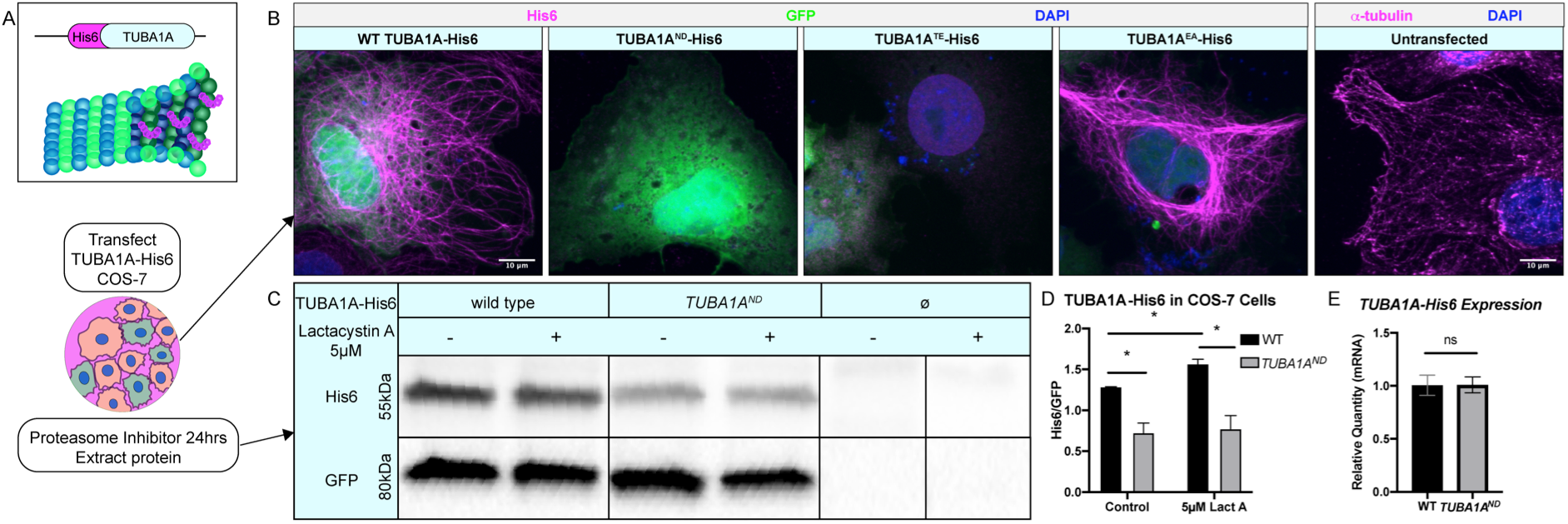
TUBA1A^ND^ impairs incorporation into cellular microtubules and reduces α-tubulin protein abundance. ***A.*** Schematic of *TUBA1A*-His6 tag and experimental design. His6 epitope tag was added to an internal loop on TUBA1A (top), and TUBA1A-His6 plasmid DNA was transfected into Cos-7 cells to label cellular microtubules in the presence or absence of proteasome inhibitor treatment (bottom). ***B.*** Images showing Cos-7 cells transfected with wild-type, *TUBA1A^ND^*, *TUBA1A*^*TE*^ (polymerization-incompetent mutant), and *TUBA1A*^*EA*^ (highly polymer stable mutant) *TUBA1A* -His6 plasmid DNA. Cells were immunolabeled with α-His6 antibodies to reveal microtubule incorporation of wild-type and mutant TUBA1A-His6 protein. TUBA1A-His6 images on the left are compared to traditional immunolabeling of α-tubulin using DM1A antibody in untransfected Cos-7 cells on right. ***C.*** Western blot for His6 in TUBA1A-His6 transfected Cos-7 cell lysates. A subset of transfected cells were treated with 5μM Lactacystin A (LactA) for 24 hours to block proteasomal degradation. His6 signal was normalized to GFP, which was expressed from the same plasmid. ***D.*** Quantification of band density for His6 western blot shown in *C*. His6 band density was normalized to GFP-expressing cells in control-treated (left), and cells treated with 5μM LactA for 24-hours (right). Data were analyzed by t test. N=3 transfections; n=3 technical replicates; p=0.04 for all comparisons marked with asterisks; p=0.83 for control vs LactA-treated *TUBA1A^ND^*. ***E.*** Bar graph representing *TUBA1A* mRNA expression in Cos-7 cells transfected with TUBA1A-His6. *TUBA1A* mRNA expression was normalized to GFP mRNA expression. Data were normalized to *TUBA1A* expression in wild-type (WT) TUBA1A-His6-transfected cells and represent 3 separate transfection experiments and 3 qRT-PCR replicates. Differences between groups were assessed by t test (p=0.97). All images were taken at 40X magnification, scale bars are 10μm. All graphs show mean of data ± SEM, *p<0.05.

His6-tagged mutant TUBA1A can be ectopically expressed in cells to evaluate the abundance and polymerization capability of mutant TUBA1A proteins. We ectopically expressed three *TUBA1A* variants in Cos-7 cells: *TUBA1A^ND^*, the mutation analogous to the *Tuba1a^ND^* mice; *TUBA1A^T349E^* (*TUBA1A^TE^)* an α-tubulin mutation shown to prevent polymerization in yeast [30]; and *TUBA1A^E255A^* (*TUBA1A^EA^*) an α-tubulin mutation which inhibits depolymerization and thus becomes locked in a polymer-bound state [28, 29](Fig. 3B). As predicted, *TUBA1A^TE^*-His6 protein was not highly detected in Cos-7 cell microtubule polymers, while *TUBA1A^EA^*-His6 protein was abundantly incorporated into cellular microtubules (Fig. 3B). In contrast to wild-type, but similar to the polymerization-incompetent *TUBA1A^TE^*-His6 protein, ectopic *TUBA1A^ND^*-His6 protein was not detected in Cos-7 cells (Fig. 3B). Western blotting of lysates from Cos-7 cells, 48-hours post-transfection revealed that *TUBA1A^ND^*-His6 protein was significantly reduced compared to wild-type *TUBA1A*-His6 (Fig. 3C, D; p=0.04), despite similar *TUBA1A* mRNA levels between wild-type and *TUBA1A^ND^* transfected Cos-7 cells (Fig. 3E; p=0.97). To evaluate if *TUBA1A^ND^* mutant protein is targeted for proteasomal degradation, we treated *TUBA1A*-His6 transfected Cos-7 cells with 5μM proteasome inhibitor, Lactacystin A (LactA [31, 32]), for 24-hours and probed for His6 abundance by western blot (Fig. 3C). Treatment with LactA significantly increased wild-type *TUBA1A*-His6 protein compared to control-treated cells, but had no effect on *TUBA1A^ND^*-His6 protein abundance (Fig. 3D; p=0.83). These results indicate that *TUBA1A^ND^* mutant protein is likely not targeted for degradation by the proteasome, but may reduce cellular TUBA1A by other mechanisms.

We next investigated if *TUBA1A^ND^* substitution impairs incorporation of TUBA1A protein into neuronal microtubule polymers (Fig. 4). Wild-type rat cortical neurons were nucleofected with *TUBA1A*-GFP, wild-type *TUBA1A*-His6, *TUBA1A^ND^*-His6 and *TUBA1A^TE^*-His6 DNA at day *in vitro* 0 (DIV 0; Fig. 4A). Following 2-days in culture (DIV 2), cells were fixed and a subset of neurons were permeabilized to remove soluble tubulin dimers (“tubulin extraction”). Extraction of soluble tubulin leaves behind only polymer-bound tubulin, enabling visualization of ectopic tubulin protein polymerization competence [37]. Neurons expressing *TUBA1A*-GFP exhibited abundant GFP signal in intact cells, but tubulin extraction revealed that GFP fusion impaired incorporation of TUBA1A into neuronal microtubule polymers (Fig. 4B). In contrast, wild-type *TUBA1A*-His6 protein was highly present in both intact and tubulin extracted neurons, demonstrating that His6-tagging does not impair polymerization ability of TUBA1A in neurons (Fig. 4C). As predicted, polymerization incompetent *TUBA1A^TE^*-His6 mutant protein was diffusely visible in intact neurons, but was absent from tubulin extracted neurons (Fig. 4C). *TUBA1A^ND^*-His6 protein was detectable at very low levels in unextracted neurons, but was not visible following tubulin extraction, indicating that *TUBA1A^ND^* impairs incorporation into neuronal microtubules (Fig. 4C). These experiments illustrate that *TUBA1A^ND^* reduces abundance of TUBA1A protein upstream of proteasomal degradation and prevents incorporation of mutant TUBA1A into cellular microtubules.

**Figure 4.**
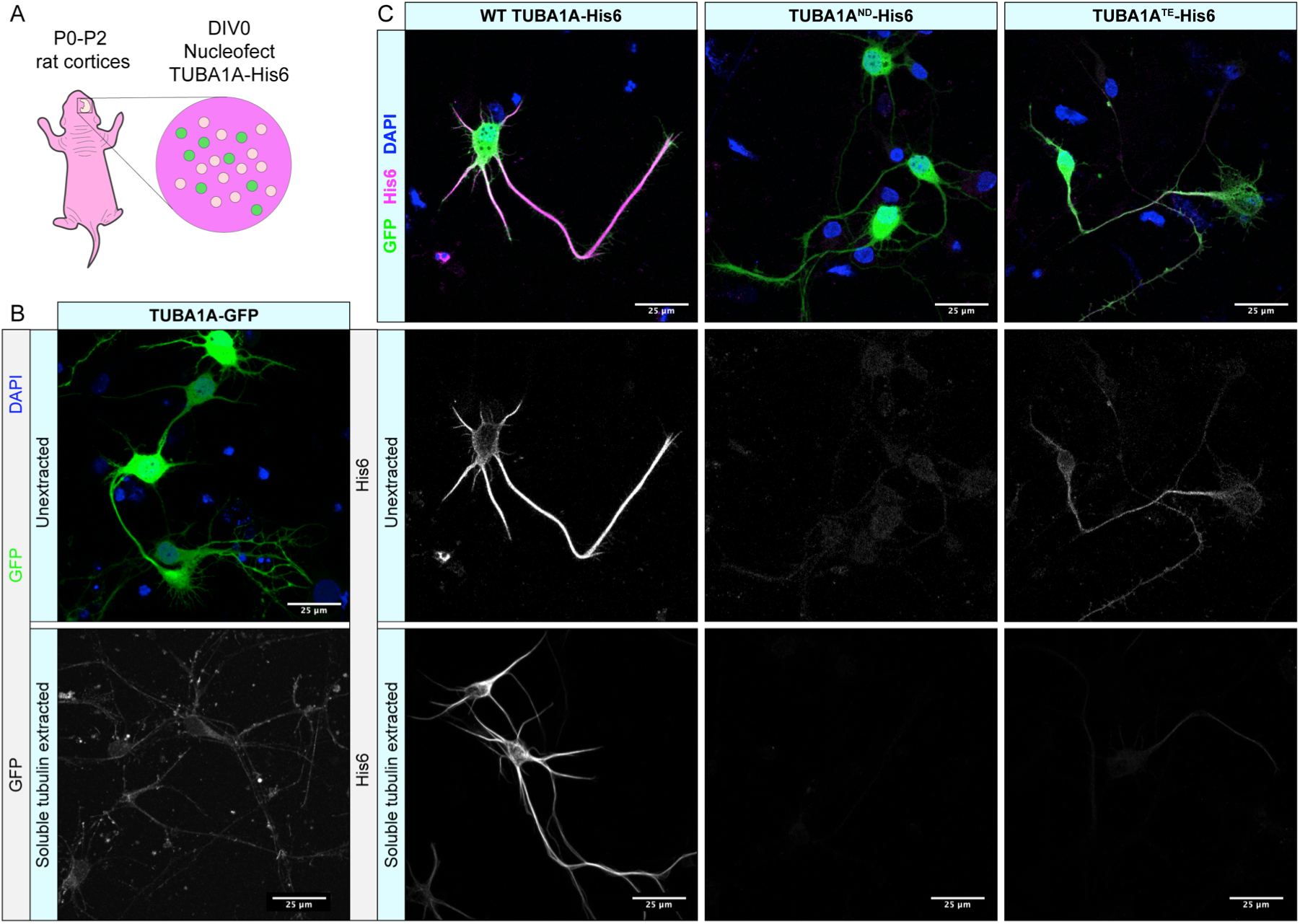
TUBA1A^ND^ α-tubulin does not incorporate into neuronal microtubule polymers. ***A.*** Schematic of cortical neuron isolation and transfection. ***B.*** Cortical rat neurons at DIV 2 transfected with *TUBA1A*-GFP. Top panel shows neurons with intracellular environment intact (unextracted), containing soluble tubulin dimers and polymerized microtubules transfected to express a TUBA1A-GFP fusion protein. Bottom panel shows neurons with soluble tubulin dimers extracted, showing only GFP-labeled TUBA1A that is incorporated into microtubule polymer. ***C.*** Rat cortical neurons at DIV 2 transfected with wild-type (WT) *TUBA1A*-His6, *TUBA1A^ND^*-His6 LOF mutation, and *TUBA1A^TE^*-His6 polymerization-incompetent mutant as a negative control. Top panels show composite image containing membrane-bound GFP (green) for confirmation of transfection, α-His6 (Magenta) and DAPI (blue) immunolabeling. Middle panels show unextracted and bottom panels show tubulin extracted DIV 2 neurons (described in *B.*), labeled with α-His6 antibodies to visualize ectopic *TUBA1A*-His6 proteins. All images were taken at 63X magnification, scale bar is 25μm.

### Reduced Tuba1a alters microtubule dynamic properties in neurons

Dynamic instability is a defining feature of microtubule polymers that is heavily influenced by a number of factors to tune microtubule network function in cells [14]. MAP-binding, PTMs, and incorporation of different tubulin isotypes can all influence microtubule dynamics [38–41]. *TUBA1A^ND^* tubulin is less abundant than wild-type and does not incorporate into neuronal microtubule polymers (Figs. 3–4). We next assessed microtubule dynamics in developing *Tuba1a^ND/+^* neurons. Wild-type and *Tuba1a^ND/+^ cortical* neurons were transfected with GFP-MACF43, a fluorescently tagged protein containing the SxIP motif bound by microtubule plus-end binding proteins (EBs), allowing for visualization of microtubule plus-end polymerization in live cells (Fig. 5A)[42]. Kymograph plots generated from time-lapse GFP-MACF43 images were analyzed for velocity, duration, and distance of GFP-MACF43 polymerization events (Fig. 5B). Previous analysis of *Tuba1a^ND/+^* GFP-MACF43 data revealed a significant reduction in the number of microtubule plus-ends per micron of neurite, which is evident in the example kymographs (Fig. 5B; [24]). Kymograph analysis showed that *Tuba1a^ND/+^* microtubules had accelerated polymerization velocity (19.67±1.1μm/min) compared to wild-type at DIV 1 (17.36±0.43 μm/min; Fig. 5C; n=688 events per genotype; p=0.0008 by Mann-Whitney test), consistent with prior data from *S. cerevisiae* [22]. Additionally, *Tuba1a^ND/+^* polymerization events covered a larger distance (Fig. 5D; n=688 events, p<0.0001) and lasted a longer amount of time (Fig. 5E; n=688 events, p<0.0001) than polymerization events in wild-type neurons. The increased polymerization rates of *Tuba1a^ND/+^* neuronal microtubules was surprising, but collectively with our previous results these data indicate that *Tuba1a^ND^* protein is not functional, and that the diminished α-tubulin pool leads to altered microtubule dynamics and changes to microtubule organization in developing neurons.

**Figure 5.**
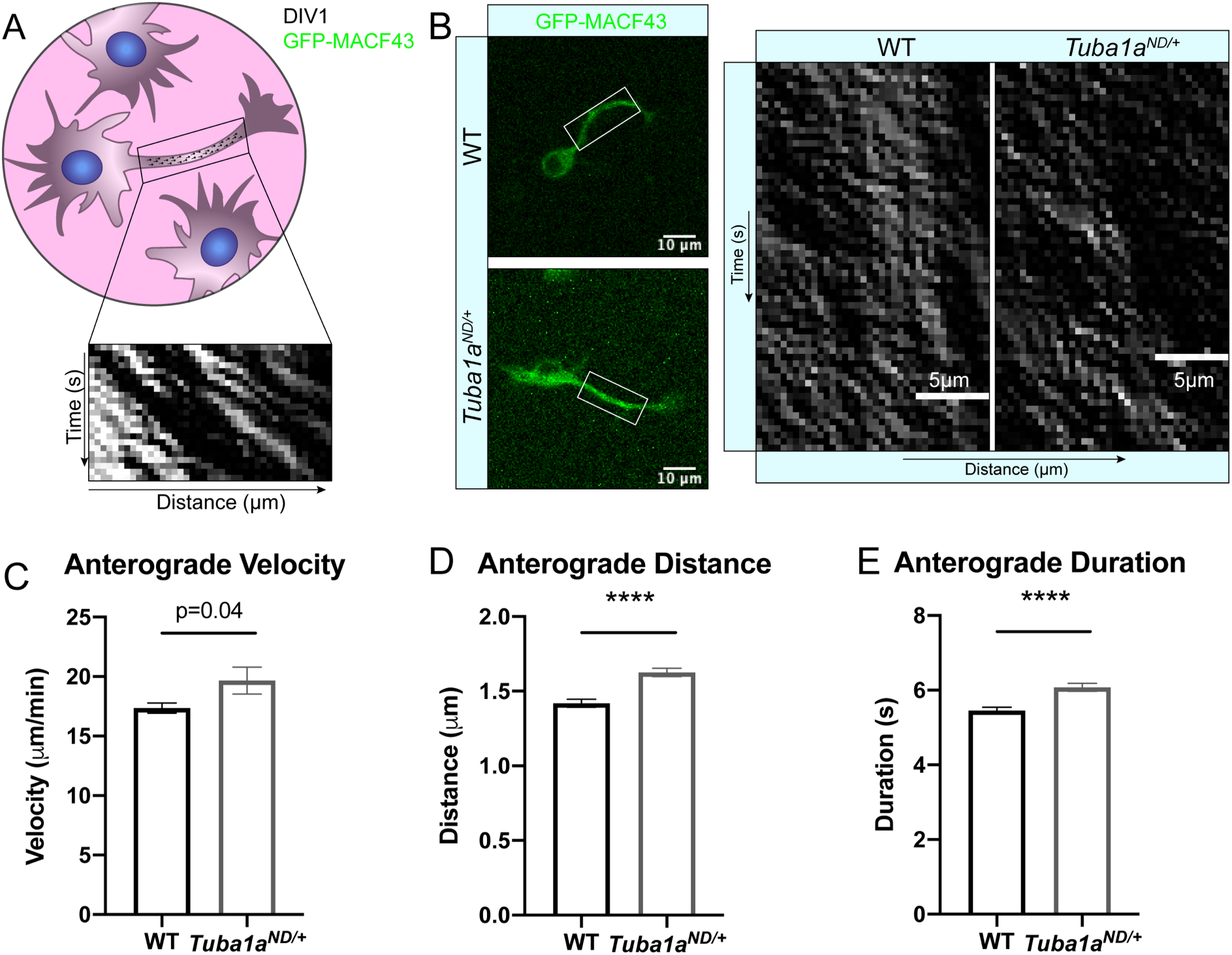
Reduced Tuba1a alters neuronal microtubule polymerization at DIV 1. ***A.*** Schematic of nucleofection and imaging experimental setup. Neurons were nucleofected with GFP-MACF43 plasmid DNA and imaged at DIV 1 for microtubule polymerization events in a neurite region of interest. ***B.*** Representative images showing GFP-MACF43 expressing wild-type (WT) and *Tuba1a*^*ND/+*^ neurons at 40X magnification (left) and representative kymographs that were generated from GFP-MACF43 images (right). Scale bars are 10μm and 5μm on neuron images and kymographs, respectively. ***C.*** Bar graph representing the average polymerization velocity for anterograde microtubule polymerization events in WT and *Tuba1a^ND/+^* cortical neurons at DIV 1 (n=688 events; p=0.0008 by Mann-Whitney test). ***D.*** Bar graph representing polymerization distance in DIV 1 WT and *Tuba1a*^*ND/+*^ cortical neurons (n=688 events, p<0.0001). ***E.*** Bar graph representing polymerization duration in DIV 1 WT and *Tuba1a*^*ND/+*^ cortical neurons (n=688 events, p<0.0001). Bars represent the mean of each group and error bars represent SEM. Differences between groups were assessed by t test, unless otherwise noted.

### *Tuba1a* is necessary for neurite extension and cytoskeletal organization in growth cone

To assess potential mechanisms by which reduced Tuba1a prevents commissural neurons from reaching their contralateral targets, we measured neurite growth rates in cultured primary cortical neurons from P0 wild-type and *Tuba1a^ND/+^* mice (Fig. 6A). We labeled primary wild-type and *Tuba1a^ND/+^* cortical neurons with a membrane-bound Myr-TdTomato to visualize growth rates in live neurons (Fig. 6A). Analysis of membrane bound Myr-TdTomato images taken one hour apart revealed that *Tuba1a^ND/+^* neurons grew at a significantly slower rate than wild-type neurons (Fig. 6B; N=3 mice, n=13 neurons; p=0.03 by Mann-Whitney test) with *Tuba1a^ND/+^* neurons growing an average of 4.67±1.15μm/hr compared to 12.60±3.12μm/hr in wild-type neurons. Additionally, measurements of the longest neurite, designated a putative ‘axon’, at DIV 3 revealed that *Tuba1a^ND/+^* neurites were significantly shorter than that of wild-type (Fig. 6C; N=3 mice, n=124 neurons; p=0.02).Together, these data show that developing neurons with reduced Tuba1a have shorter neurites and grow at a slower rate than wild-type.

**Figure 6.**
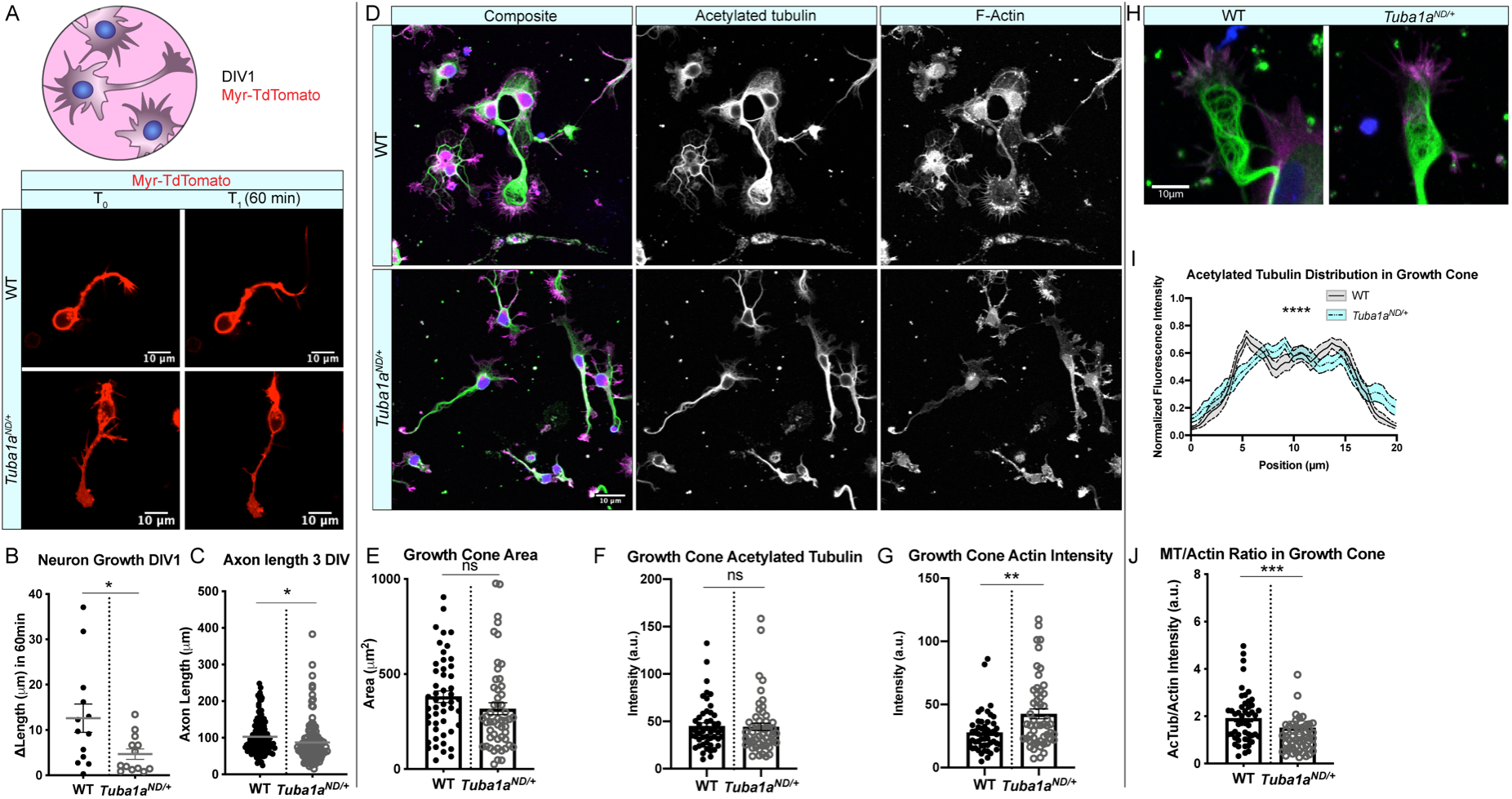
Tuba1a^ND^ impairs neurite outgrowth and alters growth cone cytoskeleton in cortical neurons. ***A.*** Schematic of experimental design and time-lapse images of membrane-labeled Myr-TdTomato neurons at DIV 1. Myr-TdTomato images were aquired 60 minutes apart to assess neuronal growth rate in wild-type (WT; top) and *Tuba1a*^*ND/+*^ (bottom) cortical neurons. ***B.*** Scatter plot of growth rates in live cortical neurons from WT and *Tuba1a*^*ND/+*^ mice at DIV 1. Data points represent individual neurons (N=3 mice, n=13 neurons; p=0.03 by Mann-Whitney test). ***C.*** Scatter plot of neurite length at DIV 3 in fixed WT and *Tuba1a*^*ND/+*^ primary cortical neurons. For each cell, the longest neurite, designated a putative ‘axon’, was measured from the cell soma to the distal neurite tip using an acetylated tubulin marker. Data points represent individual neurons (N=3 mice, n=124 neurons; p=0.02). ***D.*** Images of DIV 3 WT (top) and *Tuba1a*^*ND/+*^ (bottom) cortical neurons stained with antibodies directed against acetylated tubulin (green) and rhodamine phalloidin (F-actin, magenta). Composite images are shown (left) with single channel grayscale images for acetylated tubulin (middle) and F-actin (right). Scale bars are 10μm. ***E.*** Scatter plot of growth cone area at DIV 3 in WT and *Tuba1a*^*ND/+*^ cortical neurons. Data points represent individual growth cones (N=3 mice, n=52 growth cones; p=0.26). ***F.*** Scatter plot of acetylated tubulin intensity within the growth cone of WT and *Tuba1a*^*ND/+*^ cortical neurons at DIV 3. Data points represent individual growth cones (n=49 growth cones; p=0.89). ***G***. F-actin intensity within growth cone of WT and *Tuba1a*^*ND/+*^ cortical neurons at DIV 3. Data points represent individual growth cones (N=3 mice, n=49 growth cones; p=0.0014). ***H***. Representative images of WT and *Tuba1a*^*ND/+*^ cortical neuron growth cones showing distribution of acetylated tubulin (green) and F-actin (magenta). ***I.*** Line plot showing the average acetylated tubulin fluorescence intensity across a 20μm line scan through the central domain of the growth cone in DIV 3 WT and *Tuba1a*^*ND/+*^ neurons. Differences between WT and *Tuba1a*^*ND/+*^ growth cones were assessed by two-way ANOVA which showed a significant interaction between genotype and fluorescence intensity by position (n=20 growth cones; p<0.0001). ***J.*** Scatter plot representing the ratio of acetylated tubulin to F-actin intensity within the growth cones of DIV 3 WT and *Tuba1a*^*ND/+*^ cortical neurons. Data points represent individual growth cones (N=3 mice, n=49 growth cones; p=0.0003 by Mann-Whitney test). For all plots, lines represent mean and error bars report SEM. Differences between WT and *Tuba1a*^*ND/+*^ datasets were assessed by t test unless otherwise noted. *p<0.05; **p<0.01;

To explore the precise mechanisms by which reduced Tuba1a contributes to slowed outgrowth in *Tuba1a^ND/+^* neurons, we assessed the abundance of acetylated microtubules and filamentous actin (F-actin) in developing growth cones of wild-type and *Tuba1a^ND/+^* cortical neurons at DIV 3 (Fig. 6D). The growth cone is a dynamic developmental structure that uses the coordinated action of the actin and microtubule cytoskeleton to drive neuronal outgrowth in response to internal and external cues [43, 44]. Growth cone area was not changed in *Tuba1a^ND/+^* neurons (317.4±31.3μm^2^) compared to wild-type (380.9±30.4μm^2^; Fig. 6E; n=49 growth cones; p=0.15). We examined the amount of acetylated tubulin, a PTM associated with stable microtubules (Fig. 6F) and found there to be no difference in the overall fluorescence intensity of acetylated tubulin in *Tuba1a^ND/+^* neurons compared to wild-type (n=49 growth cones; p=0.89). In contrast, we observed a significant increase in F-actin intensity within the growth cones of *Tuba1a^ND/+^* neurons compared to wild-type (Fig. 6G; n=49 growth cones; p=0.0014). Neuronal microtubules splay out in the central, actin-dominated regions of the growth cone, but are bundled towards the peripheral domains of the growth cone [45]. To assess the degree of growth cone microtubule bundling, we next performed line scans across the widest point of DIV 3 growth cones ≥10μm (Fig. 6H). Line scans of acetylated tubulin through the growth cone revealed differences in microtubule organization between wild-type and *Tuba1a^ND/+^* neurons (Fig. 6I; n=39 growth cones; p<0.0001 between genotypes by two-way ANOVA). Specifically, we observed peaks in fluorescence, indicating bundled microtubules, at the edges of the growth cone in wild-type neurons, where acetylated tubulin was more diffuse and lacked obvious organization in *Tuba1a^ND/+^* growth cones (Fig. 6I). Intriguingly, the ratio of acetylated microtubules to F-actin in the growth cone was significantly reduced in *Tuba1a^ND/+^* neurons compared to wild-type, indicating changes to the overall growth cone cytoskeletal environment in *Tuba1a^ND/+^* neurons (Fig. 6J; n=49 growth cones per genotype; p=0.0003). Together, these data indicate that neurite growth rate is particularly sensitive to the amount of Tuba1a tubulin available, and impaired Tuba1a function leads to abnormal actin and microtubule architecture in the developing growth cone.

### *Tuba1a^ND^* neurons fail to localize critical developmental proteins to growth cone

MAPs play a crucial role in regulating neuronal microtubule function to support proper neurodevelopment. Microtubule-associated protein 1b (Map1b) promotes axon extension and is required for formation of the corpus callosum in mice [46, 47]. There was no significant deficit in the association of Map1b with microtubules from *Tuba1a^ND/+^* brain lysates compared to wild-type; in fact, *Tuba1a^ND/+^* lysates bound slightly more Map1b than wild-type (Fig. 7A, B; p=0.03). Western blot analysis of whole brain lysates from wild-type and *Tuba1a^ND/+^* mouse brains showed no difference in the total amount of Map1b protein (Fig. 7A,C; p=0.98). These data indicate *Tuba1a^ND^* does not impair Map1b’s interaction with neuronal microtubules. In developing wild-type neurons, Map1b localizes strongly to the growth cone to promote axon growth and facilitate microtubule response to guidance cues [47–51]. *Tuba1a^ND/+^* neurons contained Map1b protein, but exhibited very little Map1b fluorescence in the growth cone compared to wild-type neurons (Fig. 7D, E; n=31 growth cones; p=0.009). These data provide evidence that while the abundance of Map1b protein is unchanged by *Tuba1a^ND^*, reduced Tuba1a does not allow for correct subcellular localization of Map1b to the growth cone. Failure of *Tuba1a^ND/+^* neurons to localize Map1b to the developing growth cone provides a putative mechanism by which developing axons may fail to respond to critical guidance cues, leading to the commissural deficits observed in *Tuba1a^ND/+^* mice.

**Figure 7.**
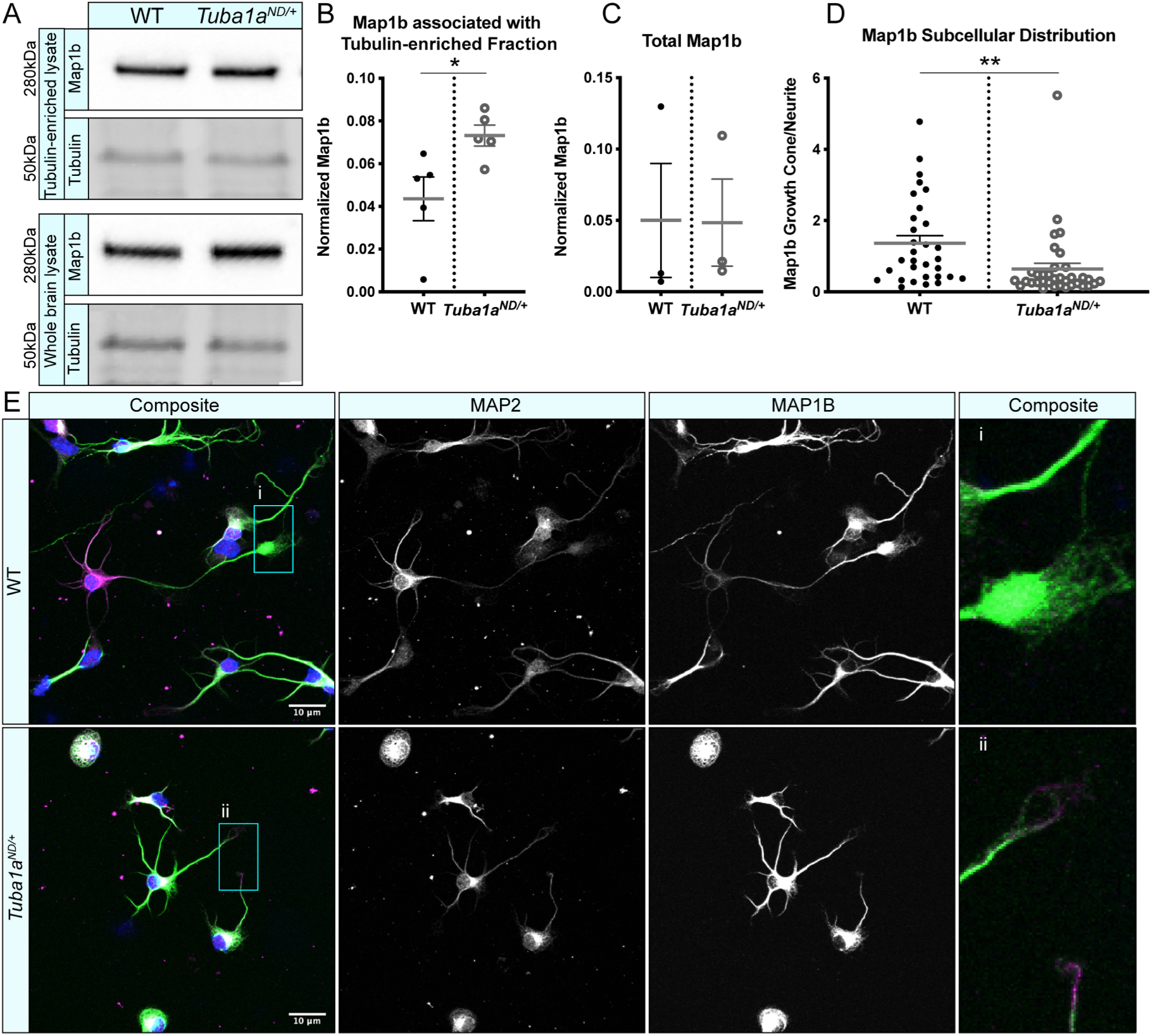
Tuba1a^ND^ neurons do not correctly localize Map1b to the developing growth cone. ***A.*** Western blots showing Map1b protein associate with a tubulin-enriched fraction from brain (top panel) and total Map1b protein in whole brain lysate (bottom panel) from wild-type (WT) and *Tuba1a*^*ND/+*^ mice. Due to the amount of protein that was loaded for Map1b western blots, antibody-stained bands for α-tubulin were oversaturated and could not be quantified, thus Map1b was normalized to the 50kDa band (presumed to be primarily tubulin) on a UV-activated stain-free blot. ***B.*** Scatter plot quantifying Map1b associated with the tubulin-enriched brain lysate, normalized to the 50kDa presumed tubulin band using stain-free western blotting. p=0.03 ***C.*** Scatter plot representing total Map1b protein in brain lysate by western blot, normalized to the total protein on a stain-free blot. p=0.98 ***D.*** Scatter plot showing the subcellular distribution of Map1b protein in WT and *Tuba1a*^*ND/+*^ cortical neurons at DIV 3. Data are represented as Map1b fluorescent signal in growth cone region divided by a region proximal to the cell body. p=0.009. ***E.*** Representative images showing altered subcellular distribution of Map1b in *Tuba1a*^*ND/+*^ (bottom) cortical neurons compared to WT (top) at DIV 3. Composite and individual channel grayscale images of MAP2 and Map1b immunocytochemistry are shown, ***i*** and ***ii*** indicate enlarged regions shown in insets. Scale bars are 10μm. Differences between groups were evaluated by t test. * p<0.05; ** p<0.01.

### Multi-faceted functions of TUBA1A during neurodevelopment

Our results support two potential models by which TUBA1A regulates neuronal microtubules to promote commissural axon pathfinding. Using our *TUBA1A*-His6 tool, we illustrated that the *TUBA1A^ND^* allele diminishes TUBA1A protein abundance in cells (Fig. 3). Here, we demonstrate that diminished Tuba1a alters microtubule dynamics and impairs neurite outgrowth *in vitro* (Figs. 5–6). Thus, reduced Tuba1a could render neurons incapable of supplying the microtubule bulk needed to drive axon outgrowth forward at a specific rate (Fig. 8A). The timing of neurodevelopment is precisely regulated, and neurons which fail to reach targets at the correct time can miss crucial developmental signaling events. We additionally show that neurons with reduced Tuba1a fail to localize a critical developmental MAP, Map1b, to the growth cone (Fig. 7). Interactions between MAPs and microtubules play a major role in adapting microtubule function in response to a changing intra- and extra-cellular developmental environment [46, 51, 52]. Therefore, neurons with reduced Tuba1a may be rendered unable to respond to extracellular guidance cues, such as Netrin1, due to failed sub-cellular localization of MAPs and other critical cargoes (Fig. 8B). Our evidence supports a multi-faceted role for TUBA1A during neurodevelopment, where it tunes microtubule dynamics and density to fuel growth and also provides stable tracks for rapid intracellular transport.

**Figure 8.**
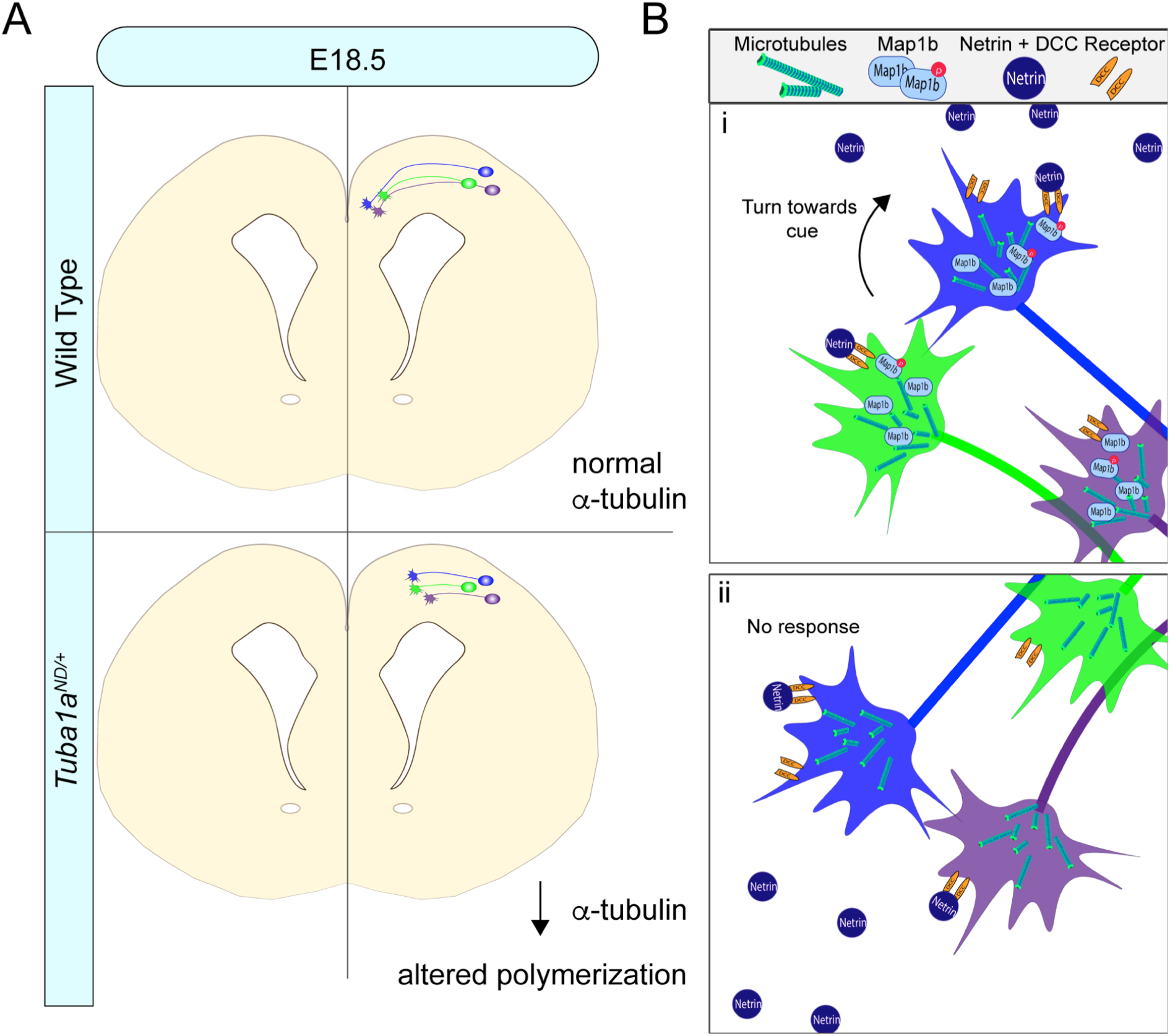
Mechanisms of Tuba1a-induced axonal pathfinding deficits. ***A.*** Schematic illustrating how reduced Tuba1a may impair ability of *Tuba1a*^*ND/+*^ axons to reach key signaling points at the correct developmental time to support proper brain formation. Wild-type axons (WT; top) reaching the midline crossing point of corpus callosum at embryonic day (E) 18.5 are compared to the potentially stunted axonal growth in *Tuba1a*^*ND/+*^ brains (bottom). The reduced density of microtubules and altered polymerization dynamics likely contribute to slower growth of developing *Tuba1a*^*ND/+*^ commissural neurons. ***B.*** Schematics illustrating potential molecular mechanisms by which Tuba1a supports navigating axons. ***i.*** Inset from *(A)*, showing WT growth cones navigating through midline corpus callosum. WT growth cones are rich with Map1b, which binds growth cone microtubules to guide cellular response to guidance molecules like Netrin. ***ii.*** Inset from *(A)* showing *Tuba1a*^*ND/+*^ growth cones, which fail to localize Map1b and may be rendered unable to mount a cytoskeletal response to extracellular cues.

## Discussion

### Studying α-tubulin isotypes *in vivo*

*TUBA1A* has long been associated with neurodevelopment due to its spatial and temporal expression as well as its role in tubulinopathies [2–4, 17–20, 53–55], but it has been historically difficult to study the contribution of a single α-tubulin isotype to microtubule network function *in vivo* due to the limited availability of isotype-specific tools. Here, we present a novel tool for studying TUBA1A protein *in vivo* without impacting native microtubule properties by introducing a hexahistidine (His6) tag into a previously identified internal loop of TUBA1A (Fig. 3) [25–27]. In this study, we provide the first evidence that TUBA1A is essential for regulating neuronal microtubule function to support long-distance targeting of central nervous system axons. Ectopic expression of *TUBA1A*-His6 protein in cells revealed that *TUBA1A^ND^* protein is approximately half as abundant as wild-type (Fig. 3D), despite similar amounts of mRNA expression in transfected cells (Fig. 3E). Based on this evidence, *TUBA1A^ND^* substitution causes targeted depletion of the mutant TUBA1A protein, but our results suggest that degradation by the proteasome is unlikely (Fig. 3D). Newly synthesized α- and β-tubulin proteins enter a complex tubulin folding pathway, where they interact with cytosolic chaperonins and tubulin-binding cofactors to become folded and assembled into tubulin heterodimers [56]. As *TUBA1A^ND^* protein is diminished compared to wild-type, but is not proteasomally degraded, we predict that mutant *TUBA1A^ND^* protein may ineffectively cycle through the tubulin folding pathway. Introduction of the ND substitution into the primary yeast α-tubulin, *Tub1* (*Tub1^ND^*) was previously shown to be lethal when combined with tubulin folding pathway mutants, providing evidence that ND substitution impairs tubulin assembly [22]. Further testing will be needed to evaluate the precise molecular mechanisms responsible for reduced *TUBA1A^ND^* protein in mammalian neurons, but it is evident that Tuba1a function is lost in the *Tuba1a^ND^* rodent model.

Neurons with reduced Tuba1a function exhibited accelerated microtubule polymerization compared to wild-type, but also demonstrated deficits in neurite extension likely due to the decreased axonal microtubule density (Figs. 5, 6; [24]). Additionally, Tuba1a-deficient microtubules were not adequate to support growth cone localization of at least one critical developmental MAP associated with commissural axon pathfinding, Map1b (Fig. 7). We show that Tuba1a-rich microtubules promote axon outgrowth and pathfinding, and reduced Tuba1a was not sufficient to form forebrain commissures in *Tuba1a^ND/+^* mice (Figs. 1–2). Collectively, these data support the conclusion that reduced Tuba1a during neurodevelopment is adequate for cortical neuron migration [22], but does not allow for sufficient microtubule function to properly localize proteins to the growth cone for axon guidance. Thus, long range axon guidance may be exquisitely sensitive to α-tubulin levels and microtubule structure.

Studying individual α-tubulin isotypes in neurons has been historically arduous as the high degree of amino acid sequence similarity between α-tubulin isotypes has prevented generation of a TUBA1A-specific antibody and has made genetically targeting a single α-tubulin gene challenging. The abundance of clinically identified mutations to *TUBA1A* provide strong evidence that *TUBA1A* is a major player in both tubulinopathy and typical neurodevelopment; however, the lack of available tools to study TUBA1A *in vivo* has prevented researchers from understanding precisely how *TUBA1A* contributes to neurodevelopment. As such, previous studies of tubulinopathy mutations have relied heavily on mRNA analysis and indirect methods of evaluating TUBA1A function. Here we introduce an important advancement in the study of TUBA1A protein *in vivo*, by harnessing a previously-identified internal loop within TUBA1A that tolerates addition of small epitope tags without impacting TUBA1A incorporation or dynamics [26]. Our internal *TUBA1A-His6* construct marks an important advancement for the study of tubulinopathies and neuronal α-tubulin as a whole. *TUBA1A-His6* was readily expressed in both Cos-7 cells and neurons and was able to incorporate into microtubule polymers, unlike *TUBA1A* containing a GFP fusion that prohibited incorporation into neuronal microtubules (Fig. 4). Additionally, we successfully used this epitope-tagged TUBA1A to model mutant tubulin behavior *in vitro* (Figs. 3–4. Overall, this tool provides an important advancement in the study of α-tubulin protein *in vivo*, and makes interrogating the function of specific α-tubulin isotypes accessible to more researchers.

### Tuba1a influences microtubule density, dynamics, and function in cells

The microtubule cytoskeleton supports a wide range of different cellular functions in different cell types, ranging from facilitating chromosome segregation during mitosis to forming dynamic and motile structures like the neuronal growth cone. Understanding how different cells use the same basic building blocks to create vastly different microtubule-based structures is a major question in microtubule biology. Many different mechanisms have been identified through which cells can regulate microtubule network properties and overall function. We show that a mutation which reduced Tuba1a incorporation into cellular microtubules accelerates microtubule polymerization rates (Fig. 5). Alterations to tubulin isotype composition were previously shown to change microtubule polymerization dynamics when the analogous mutation was made in yeast, *Tub1^ND^*. A similar acceleration of microtubule polymerization was observed in *Tub1^ND^* yeast mutants and was likely caused by a shift in the α-tubulin isotype ratio at the protein level [22]. We propose that this polymerization rate increase is not due to a reduction of available cellular tubulin, but instead is due to a change in the ratio of tubulin isotypes available for microtubule growth. This result is also supported by recent evidence showing that increased incorporation of Tuba1a tubulin, with subsequent decreased incorporation of alternative tubulin isotypes, slowed microtubule polymerization rates in *in vitro* reconstituted microtubules [40]. These results are fitting with the “tubulin code” model, which proposes that incorporation of different tubulin isotypes can modify microtubule network behavior [14, 57]. Importantly, previous work has shown that local changes to growth cone microtubule dynamics facilitate growth cone turning in response to extracellular cues [11, 13, 43, 45, 58, 59]. Thus, dysregulation of growth cone microtubule dynamics, as was observed in *Tuba1a^ND/+^* cortical neurons, could diminish the ability of developing neurons to appropriately interact with their environment. Together, these data support the conclusion that incorporation of Tuba1a α-tubulin tunes neuronal microtubule polymerization rates to support neurodevelopmental processes.

While the abundance of acetylated tubulin was not significantly different between *Tuba1a^ND/+^* and wild-type growth cones, the distribution of acetylated microtubules was different by genotype (Fig. 6). Differences in distribution of acetylated tubulin within the growth cone likely reflects altered microtubule organization (Fig. 6). Acetylation is a microtubule PTM that is associated with stable microtubule populations and as such is sparse in dynamic structures like growth cones [60–65]. However, microtubule acetylation can be induced in growth cones following contact with extracellular matrix proteins and was shown to promote cortical neuron migration *in vivo* and suppress axon branching *in vitro*, demonstrating a clear role for this PTM in development [66–68]. Tubulin PTMs, like acetylation, have been shown to impact MAP-binding affinity and function, providing a clear mechanism by which changing the PTM landscape of microtubules could alter neuronal microtubule function [41, 57, 65, 69–72]. Thus, any changes to the organization or distribution of acetylated microtubules in the growth cone could impact the ability of developing neurons to appropriately navigate their environment and establish correct synaptic targets.

*Tuba1a^ND/+^* growth cones showed a significant increase in F-actin signal compared to wild-type, causing an overall shift in the growth cone microtubule-actin balance (Fig. 6). It is well established that interplay between the actin and microtubule cytoskeleton drives growth cone movements in developing neurons [43, 44, 73, 74]. Growth cone microtubule polymerization has been shown to induce F-actin assembly, and coordination of actin and microtubules is regulated by interactions with MAPs to drive appropriate growth cone response [75–78]. As actin and microtubules are tightly regulated within the growth cone, it is reasonable to assume that mutations which disrupt microtubule function, like *Tuba1a^ND^*, likely also impact the actin cytoskeleton of developing neurons. In *Tuba1a^ND/+^* neurons, the actin cytoskeleton may occupy increased growth cone territory as the result of microtubule deficiencies, but additional testing of actin-response in developing *Tuba1a^ND/+^* neurons is needed to assess whether the increase in growth cone actin has any functional consequences. Further, we showed that *Tuba1a^ND/+^* neurons do not effectively localize at least one developmental MAP, Map1b, to the growth cone (Fig. 7). Map1b acts downstream of several important developmental signaling pathways to regulate function of both actin and microtubules within the growth cone, and dysregulation of this or other MAPs could therefore impact multiple cytoskeletal components [50, 51, 79]. The mechanisms by which *Tuba1a^ND^* induces changes to the growth cone actin cytoskeleton remain to be explored, but could reveal important insights on how microtubules and actin are coordinately regulated to support growth cone navigation.

### Models of Tuba1a-dysfunction reveal potential roles in developmental signaling

*Tuba1a^ND/+^* neuronal microtubules were not sufficient to support growth cone localization of Map1b (Fig. 7). Microtubules are the tracks upon which intracellular cargo transport occurs in neurons. Here we present evidence that impaired Tuba1a function in neurons causes aberrant localization of Map1b (Fig. 7). Map1b mRNA is a known target of the mRNA transport protein FMRP and is locally translated within developing neurons [80, 81]. As we previously demonstrated that intracellular transport is impaired in developing *Tuba1a^ND/+^* neurons [24], this provides a putative model by which reduced Tuba1a could lead to altered localization of developmental MAPs. Intracellular transport is a crucial function of neuronal microtubules throughout life; however, microtubule-based transport has been shown to be essential during neurodevelopment [12, 82–87]. Correct localization of developmental MAPs, mRNAs and organelles are crucial for cytoskeletal response to extracellular guidance cues [84, 86, 88–90]. In particular, Map1b is required for neuronal response to the guidance cue, Netrin1, a key player in commissural formation [91–96]. We showed that neuronal microtubules with reduced Tuba1a do not support neuronal growth to the same degree as wild-type microtubules, causing shorter neurite length and slower growth rates *in vitro* (Fig. 6). The timing of developmental processes is crucial for effective signal transduction and supports the formation of appropriate synaptic contacts [97, 98]. Collectively, the data presented in this study support two potential models by which *Tuba1a^ND^* neuronal microtubules fail to support proper neurodevelopment. The first model proposes that the timing of axon extension during development is crucial for effective axon guidance, as neurons with reduced Tuba1a exhibit impaired neurite extension (Fig. 6). If neurons lacking TUBA1A are not reaching the correct location at the correct time, it is possible that neurons will fail to receive key developmental signals (Fig. 8).The second model posits that neurons lacking functional TUBA1A do not have adequate intracellular transport to support localization of critical developmental proteins (Fig. 8). Inappropriate protein localization during critical points in axon extension and guidance could render neurons unable to respond to incoming guidance cues, as the machinery required to induce microtubule response to extracellular cues is absent. Importantly, these models are not mutually exclusive, as we demonstrated that TUBA1A is crucial for both developmental protein localization and neuron outgrowth. The extent to which these processes contribute to the overall deficit in commissural axon guidance remains to be explored in future studies.

Understanding the mechanisms by which microtubules contribute to discrete aspects of neurodevelopment is an active area of research. Human neurodevelopmental disorders that impact microtubule function, such as tubulinopathies, demonstrate that microtubules are critical for proper neurodevelopment to occur. Tubulinopathy patients exhibit severe, sometimes lethal, brain malformations that frequently impact multiple neurodevelopmental processes, including neuronal survival, migration and axon extension [3, 6–8, 53, 99–101]. The range of phenotypes exhibited by tubulinopathy patients have made it challenging for scientists to pinpoint specific aspects of neuronal function that are reliant on TUBA1A tubulin. In this way, mutations such as the *Tuba1a^ND^* variant whose severity can be tuned according to gene dosage can be used as important tools to interrogate the requirement for Tuba1a in discrete aspects of neurodevelopment. Importantly, though tubulinopathy patients exhibit a range of brain phenotypes, commissural abnormalities such as agenesis of the corpus callosum, are one of the most commonly reported features of this disease [1–4, 20]. Cortical malformations and neuronal migration errors are also common features of *TUBA1A* tubulinopathies; however, it has thus far been unclear as to whether commissural deficits occur as a primary or secondary consequence of TUBA1A dysfunction. In this study, we provide evidence that neurons deficient in Tuba1a fail to properly navigate to meet contralateral binding partners. These data demonstrate that TUBA1A is required for forebrain commissural formation, independent of its role in neuronal survival or migration. The insights presented in this manuscript expand upon the currently known role for *TUBA1A* in neurodevelopment, and advance the study of tubulinopathy by presenting specific mechanisms by which TUBA1A supports neurodevelopment.

